# A dynamic RNA hub facilitates activation induced cytidine deaminase recruitment to the immunoglobulin heavy chain locus

**DOI:** 10.64898/2026.01.19.700375

**Authors:** Mariia Mikhova, Mackenzie Kapanka, Li Han, Jens C. Schmidt, Kefei Yu

## Abstract

Activation-induced cytidine deaminase (AID) converts cytosines to uracils within the actively transcribed switch regions to initiate DNA repair and formation of DNA double strand breaks required for immunoglobulin class switch recombination (CSR). How AID specifically targets switch regions remains a key unanswered question. Using a multimodal live cell single molecule imaging approach, we demonstrate that intronic switch regions promote robust transcription by enhancing polymerase loading and persistent transcriptional bursts, resulting in the formation of a dynamic RNA hub consisting of numerous nascent switch transcripts simultaneously tethered to the *IgH* locus. We further demonstrate that AID interacts with switch region RNA *in vivo*, and that this interaction is required for recruitment of AID to the *IgH* locus. Together, our findings show that the RNA hub formed by nascent switch region transcripts may be part of a “class switch recombination center” and drives the specific recruitment of AID to the *IgH* locus to initiate CSR.

## RESULTS

In mammals, B lymphocytes produce several classes of immunoglobulins (Ig), each tailored to mediate distinct effector functions for effective adaptive immune responses. The Ig class (or isotype) is defined by the constant (C) region of the Ig heavy (H) chain polypeptide, while antigen specificity is conferred by the variable (V) regions of both heavy and light chains. In its germline configuration, the mouse *IgH* locus contains 7 constant regions in tandem (10 in human) spread out over ∼160 kilobases (∼250 kb in human) (Fig. 1A) (Chaudhuri and Alt, 2004; Feng et al., 2020; Yu and Lieber, 2019; Leeman-Neill et al., 2024; Methot and Di Noia, 2017). Naïve B cells co-express IgM and IgD encoded by Cµ and Cδ, respectively, via alternative RNA splicing. Expression of a distal constant region (e.g., γ, ε, α) requires induction and repair of DSBs in switch regions preceding the constant region exons, resulting in an intrachromosomal deletion known as class switch recombination (CSR, Fig. 1A) (Chaudhuri and Alt, 2004; Feng et al., 2020; Yu and Lieber, 2019; Leeman-Neill et al., 2024; Methot and Di Noia, 2017).

**Figure 1.**
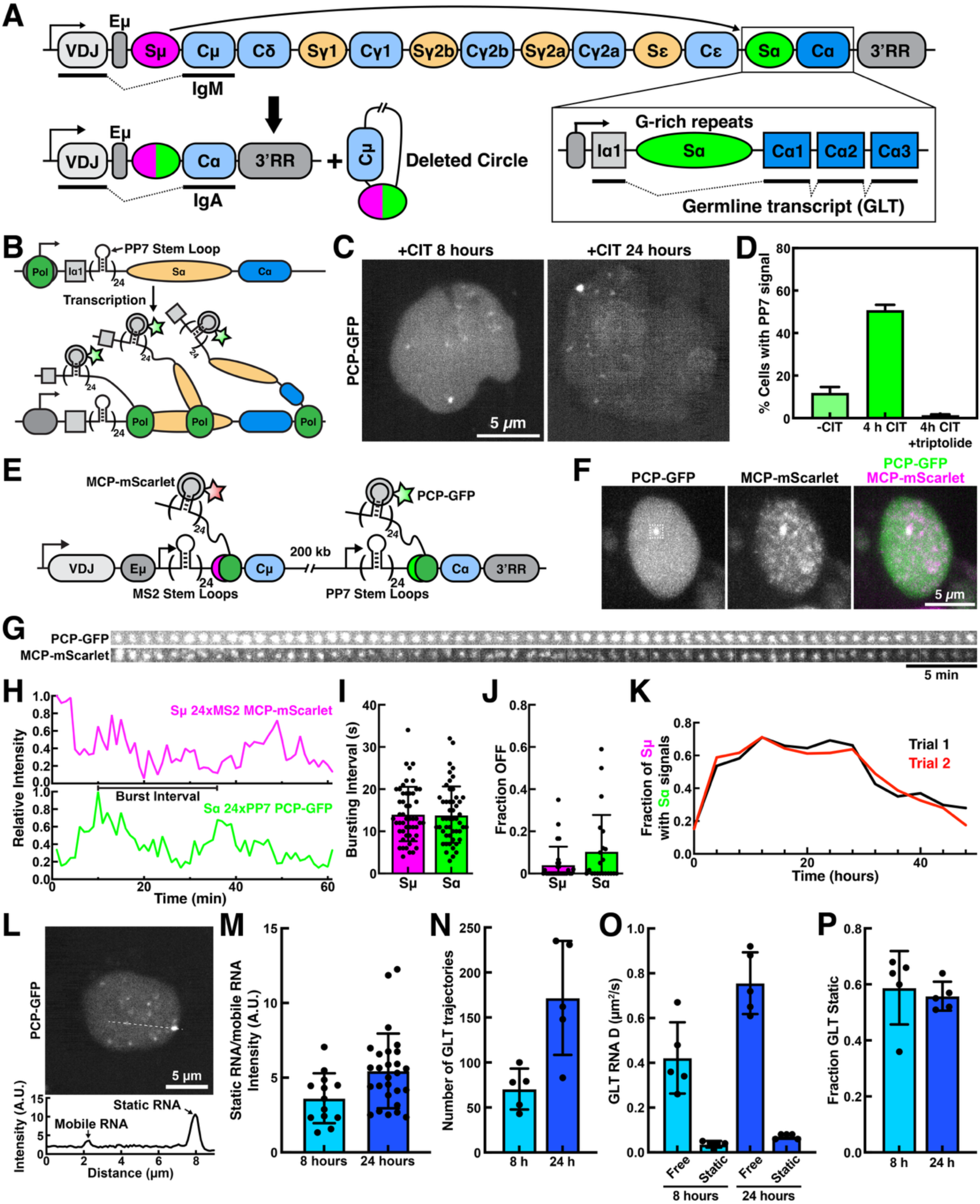
Analysis of germline transcription in living cells. **(A)** Model of class switch recombination. **(B)** Experimental design for detection of germline transcripts of the Sα region using a 24xPP7 stem loop array and PCP-GFP. **(C)** Images of living CH12 cells with the 24xPP7 array integrated upstream of the Sα switch region expressing PCP-GFP. **(D)** Quantification of the fraction of cells containing static transcriptional PCP-GFP foci after stimulation with CIT or stimulation with CIT and treatment with triptolide (N = 2 biological replicates, >50 cells per replicate, Mean±SD). **(E)** Experimental design for simultaneous detection of the transcription of the Sµ and Sα regions of the *IgH* locus using 24xMS2 and 24xPP7 stem loop arrays with MCP-mScarlet and PCP-GFP. **(F)** Single frame of a living CH12 cells containing the 24xMS2 and 24xPP7 stem loop arrays depicted in (E) expressing MCP-mScarlet and PCP-GFP imaged every minute for 60 minutes. **(G-H)** PCP-GFP and MCP-mScarlet signals **(G)** and their quantification **(H)** from every frame of a movie acquired every minute for 60 minutes of the cell depicted in (E). Also see Movie S3. **(I-J)** Quantification of the **(I)** transcriptional burst frequency and **(J)** fraction of time that no transcriptional foci are observed from the Sµ and Sα switch regions (n = 28 and 22 cells). **(K)** Quantification of the fraction of cells with a detectable Sµ transcriptional signal that co-localized with an Sα signal (n > 20 cells per timepoint). **(L)** Image of a living CH12 cells with the 24xPP7 array integrated upstream of the Sα switch region expressing PCP-GFP (top) and line scan intensity analysis of the PCP-GFP signal along the dashed line (bottom). **(M)** Quantification of the intensity of static, nascent RNA signal relative to the signal of mobile RNA molecules. **(N-P)** Single particle tracking analysis of PCP-GFP signals not localized at the *IgH* locus showing **(N)** the number of trajectories with at least 3 localizations, **(O)** the diffusion coefficients of mobile and static PCP-GFP particles, and **(P)** the fraction static RNA molecules (N = 5 biological replicates, >20 cells per replicated, Mean±SD).

CSR is initiated by activation-induced cytidine deaminase (AID) which deaminates cytosines to uracils on both strands of actively transcribed switch regions (Kim et al., 2020; Xue et al., 2006). The uracils generated by AID are processed into DSBs by the base excision and mismatch repair factors (Imai et al., 2003; Rada et al., 2002; Rada et al., 2004). AID deaminates single-stranded (ss) DNA and its activity *in vivo* requires transcription. However, how AID specifically targets actively transcribed switch regions during CSR remains a critical unaddressed question in immunology (Liu et al., 2008; Yamane et al., 2011; Álvarez-Prado et al., 2018; Meng et al., 2014; Pefanis et al., 2014; Qian et al., 2014). Switch regions are long (2-10 kb), repetitive sequences with a G-rich non-template strand (Yu and Lieber, 2003). Switch region transcription (termed germline transcription) initiates from a non-coding exon (termed “I” exon) upstream of the switch region and terminates downstream of the constant region, producing a G-rich RNA transcript that is spliced to excise the switch region in the first intron (Fig. 1A) (Yu and Lieber, 2019; Yu and Lieber, 2003). Due to these sequence features, switch region transcription can result in the formation of an R-loop; a secondary structure that consists of an DNA/RNA hybrid between the template DNA and the RNA transcript displacing the non-template DNA strand (Yu et al., 2003). R-loop formation provides a simple explanation for how single-stranded DNA on the non-template strand is exposed and targeted by AID. In addition, the displaced non-template DNA strand or the RNA transcript may fold into G-quadruplex (G4) structures (Duquette et al., 2004; Qiao et al., 2017; Zheng et al., 2015). Biochemical studies have demonstrated that AID associates with branched structures (e.g. protruding tails of G4) with much higher affinity (Qiao et al., 2017; Zheng et al., 2015) than ssDNA. Because AID binds RNA and DNA with similar affinities (Qiao et al., 2017), it is possible that RNA, particularly the germline transcript, may directly contribute to AID targeting and regulation in cells. Indeed, it has been proposed that the debranched intron containing the switch region can associate with AID and target it to the *IgH* locus in trans and in a sequence-specific manner (Zheng et al., 2015). However, how released RNA transcripts can re-associate with the *IgH* locus was not defined. Neither the R-loop nor G4 structures can adequately explain why certain AT-rich sequences can substitute for an endogenous switch region and support fairly robust CSR in mouse B cells (Zarrin et al., 2004). Recently, chromatin loop extrusion was shown to be a key mechanism that controls CSR (Zhang et al., 2021). This model postulated the existence of a CSR recombination center (CSRC) that consists of the donor Sµ switch region and multiple transcription enhancers and that access to the CSRC via cohesin mediated loop extrusion is essential for the deamination of the acceptor switch region by AID. However, how AID is recruited to the CSRC has not been defined.

A key barrier to elucidating the molecular mechanisms of CSR has been the inability to track germline transcription and AID recruitment to the *IgH* locus in real-time in living cells. To overcome this challenge, we have developed a single-molecule live-cell imaging approach that allows direct visualization of individual AID molecules and switch region transcripts in living B cells undergoing cytokine-induced CSR. Our work demonstrates that high levels of rapid switch region transcription led to the formation of a dynamic RNA hub at the *IgH* locus. The RNA hub, containing many switch region transcripts with high affinity binding sites for AID, serves as a landing platform to recruit AID and facilitate cytosine deamination specifically at the *IgH* locus.

## RESULTS

### Germline transcription seeds the formation of a dynamic RNA hub

To analyze germline transcription in living cells, we integrated 24 copies of the PP7 stem loop upstream of the Sα switch region of the productive *IgH* allele (i.e. rearranged VDJ) in the CH12F3.2A mouse B-cell line (hereafter abbreviated as CH12) containing a single *IgH allele* (Fig. 1B, Fig. S1A,B). The PP7-stem loop RNA forms as a high affinity binding site for the PP7 bacteriophage coat protein (PCP) fused to super-folder GFP (referred to as GFP), allowing visualization of nascent and released RNA transcripts in living cells (Fig. 1B).

We confirmed that a transcript containing the 24xPP7 stem loops was produced and did not impact the use of alternative splice donor sites within the Sα germline transcript (Fig. S1A, S1B). Furthermore, we verified that introduction of the PP7 array and expression of PCP-GFP did not affect CSR efficiency (Fig. S1C,D).

Induction of CSR using anti-CD40 antibody, interleukin-4 and transforming growth factor β1 (CIT) resulted in the transcription dependent formation of a nascent RNA signal in a single location in the nucleus of CH12 cells (Fig. 1C,D, Movie S1-2). To determine whether this observation was generalizable to other switch regions, we integrated an additional 24xMS2 stem loop array upstream of the Sµ switch region (Fig. 1E, Fig. S1E), which did not impact CSR efficiency (Fig. S1F), and expressed the MS2 coat protein fused to the mScarlet fluorescent protein (MCP-mScarlet) and PCP-GFP (Fig. 1E). Using this approach, we detected nascent Sµ and Sα RNA signals co-localized in the nuclei of living CH12 cells (Fig. 1F, Movie S3). We observed fluctuations of intensity of the nascent RNA signals from the Sα and Sµ regions, consistent with transcriptional bursting, which occurred with similar frequencies for both switch regions (Fig. 1G-I, Fig. S1G, Movie S3). In most cases, the nascent RNA signal did not disappear in between transcriptional bursts (Fig. 1J), suggesting that germline transcription is either stalled or reinitiated prior to termination of the previous burst, resulting in the continuous association of nascent germline transcripts with the *IgH* locus (Fig. 1G,H,J, Fig. S1G, Movie S3).

CH12 cells undergo Sµ to Sα CSR, deleting the 24xPP7 stem loop array upstream of the Sα switch region while retaining the 24xMS2 loop array upstream of the Sµ switch region (Fig. 1A). To determine the overall kinetics of CSR at the single cell level, we quantified the fraction of cells with detectable Sµ transcription signal (MCP-mScarlet) that co-localized with an Sα transcriptional signal (PCP-GFP) over a period of 48 hours (Fig. 1K). The fraction of Sµ containing cells with a co-localized Sα signal rapidly increased from 20% prior to simulation to 60% 4 hours after CIT addition, remained at ∼60% until 28 hours before continuously dropping down to ∼20% from 32-40 hours (Fig. 1K), suggesting that CSR took ∼24 hours to complete, which is consistent with the CSR kinetics detected by the expression of IgA on the cell surface of CH12 cells (Han and Yu, 2008; Kinoshita et al., 2001).

In addition to the static nascent RNA signal, we also observed less intense mobile RNA signals in the nucleus derived from both the Sµ and Sα (Fig. 1 C,F, Movie S1-2, S4). To exclude the possibility that the mobile RNA molecules were retained in the nucleus due to the nuclear localization signal (NLS) on the bound PCP-GFP molecules, we confirmed that these mobile transcripts were localized in the nucleus in cells expressing PCP-GFP without an NLS (Fig. S1I, Movie S5). The intensity of the nascent RNA foci was on average 4-5 fold (up to 12 fold) higher than the mobile RNA molecules (Fig. 1L,M), suggesting that on average 4-5 Sα switch region transcripts are simultaneously associated with the *IgH* locus. Furthermore, we observed mobile RNA molecules emanating from and reassociating with nascent RNA foci (Movie S6), indicating that the bright RNA signals may contain both tethered and untethered switch region transcripts. Single particle tracking of the mobile RNA molecules demonstrated that the number of germline transcripts released from the *IgH* locus increased over time (Fig. 1N, Fig. S1J, Movie S7). In addition, this analysis revealed that germline transcripts exist in two states, freely diffusing through the nucleus or immobile (Fig. 1O,P). Immobile particles account for 50-60% of the germline transcripts (Fig. 1P) and could represent RNA molecules associated with chromatin or RNA rich nuclear sub-compartments (i.e. nucleoli, Cajal bodies, nuclear speckles, etc.).

Together, these experiments demonstrate that germline transcription results in almost continuous association of nascent RNA with the *IgH* locus. In addition, mobile RNA molecules containing the switch region sequences accumulate in the nucleus over the course of CSR and can potentially reassociate with nascent RNA foci.

### Transcription rapidly proceeds through the Sα switch region

The persistent association of nascent germline transcripts with the *IgH* locus could be the result of transcriptional stalling within the switch region (Rajagopal et al., 2009; Wang et al., 2009). Alternatively, frequent transcriptional bursts without extended off-time could also result in continuous presence of nascent transcripts bound to the *IgH* locus. To analyze how transcription proceeds through the Sα switch region, we inserted a 24xMS2 stem loop array downstream of the Sα switch region in addition to the 24xPP7 array inserted upstream of Sα (Fig. 2A, Fig. S2A). These insertions only moderately affect CSR efficiency (Fig. S2B). Using this approach we observed bursts of PCP-GFP signal rapidly followed by the MCP-mScarlet signal, consistent with processive transcription through the switch region at a median rate of ∼2.5 kb per minute (Fig. 2B-E, Movie S8). These results demonstrate that transcription rapidly proceeds through the Sα switch region and are inconsistent with prolonged polymerase stalling within this repetitive sequence as previously suggested (Rajagopal et al., 2009; Wang et al., 2009).

**Figure 2.**
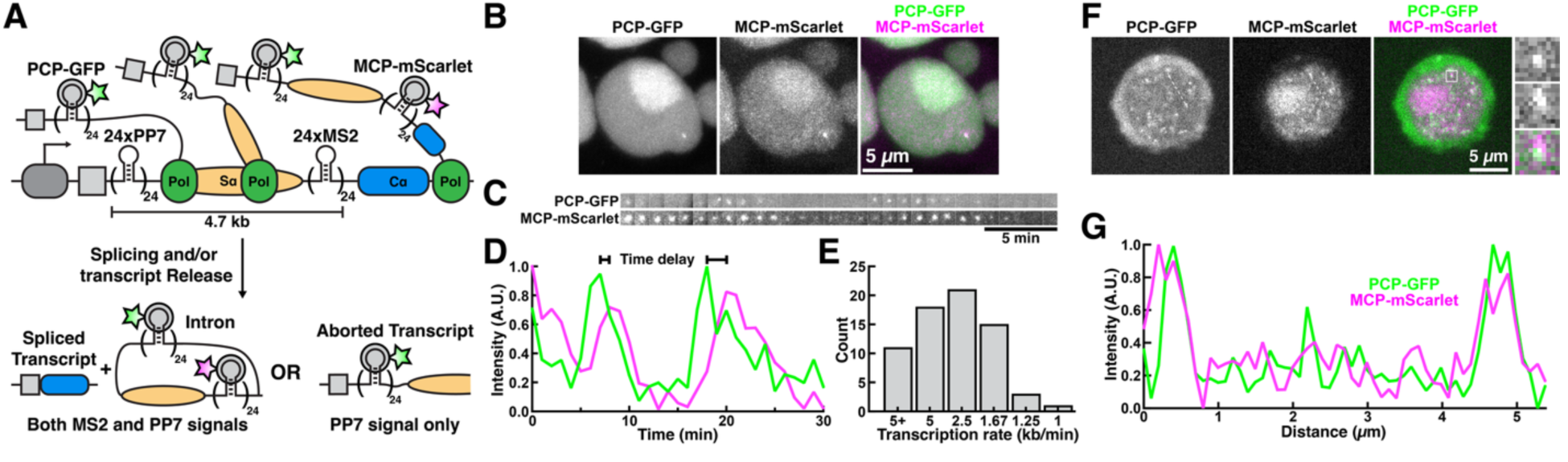
Fast transcriptional elongation across the Sα switch region. **(A)** Experimental design to analyze progression of transcription through the Sα switch region. **(B)** Images of living CH12 cells expressing PCP-GFP and MCP-mScarlet containing 24xPP7 and 24xMS2 stem loop arrays up- and down-stream of the Sα switch region, respectively. **(C-D)** Images and quantification of PCP-GFP and MCP-mScarlet signals in living CH12 cells containing 24xPP7 and 24xMS2 stem loop arrays up- and down-stream of the Sα switch region, respectively. **(E)** Quantification of the transcription rate through the Sα switch region (N=69 burst pairs over 3 biological replicates). **(F)** Full images, magnified foci (white box), and **(G)** line scan quantification (dashed line) of mobile PCP-GFP and MCP-mScarlet labeled RNA molecule intensity.

### Intronic RNA containing the Sα switch region accumulates in the nucleus of CH12 cells

The mobile nuclear germline transcripts could represent either spliced-out intronic sequences or transcripts that were terminated within the Sα switch region. To distinguish between these models, we took advantage of our engineered cell line containing 24xPP7 and 24xMS2 stem loop arrays flanking the Sα switch region (Fig. 2A). Spliced out intronic sequences would include both stem loop arrays, while transcripts aborted within the switch region would only contain the 24xPP7 stem loop array. Dual-color imaging revealed that all mobile RNA molecules are marked by both PCP-GFP and MCP-mScarlet (Fig. 2F,G, Movie S9). This observation demonstrates that spliced-out intronic RNA molecules containing the Sα switch region sequence, rather than transcripts aborted within the switch region, accumulate in the nucleus of CH12 cells during CSR. This result was unexpected since most intronic RNAs are rapidly debranched and degraded.

### Switch region sequences increase polymerase loading in cis

To address the question whether the persistent presence of nascent RNA molecules is a unique feature of switch region transcription, we inserted the 24xPP7 stem loop array into the first intron of an engineered *AID* locus containing LoxP sites for recombination mediated cassette exchange (RMCE) (Fig. 3A). The *AID* promoter is highly active upon CIT stimulation based on previous Gro-seq analysis (Zhang et al., 2019). However, compared to germline transcription at the Sα switch region, we observed long periods without active transcription at the *AID* locus, while the time between transcriptional bursts was slightly shorter at the *AID* locus (Fig. 3B-D, Fig. S3A, Movie S10). Furthermore, the peak intensity of the transcriptional bursts at the *AID* locus was ∼50% lower relative to transcriptional signals at the Sα switch region (Fig. 3E), suggesting that fewer RNA polymerases are loaded during each transcriptional burst at the *AID* locus.

**Figure 3.**
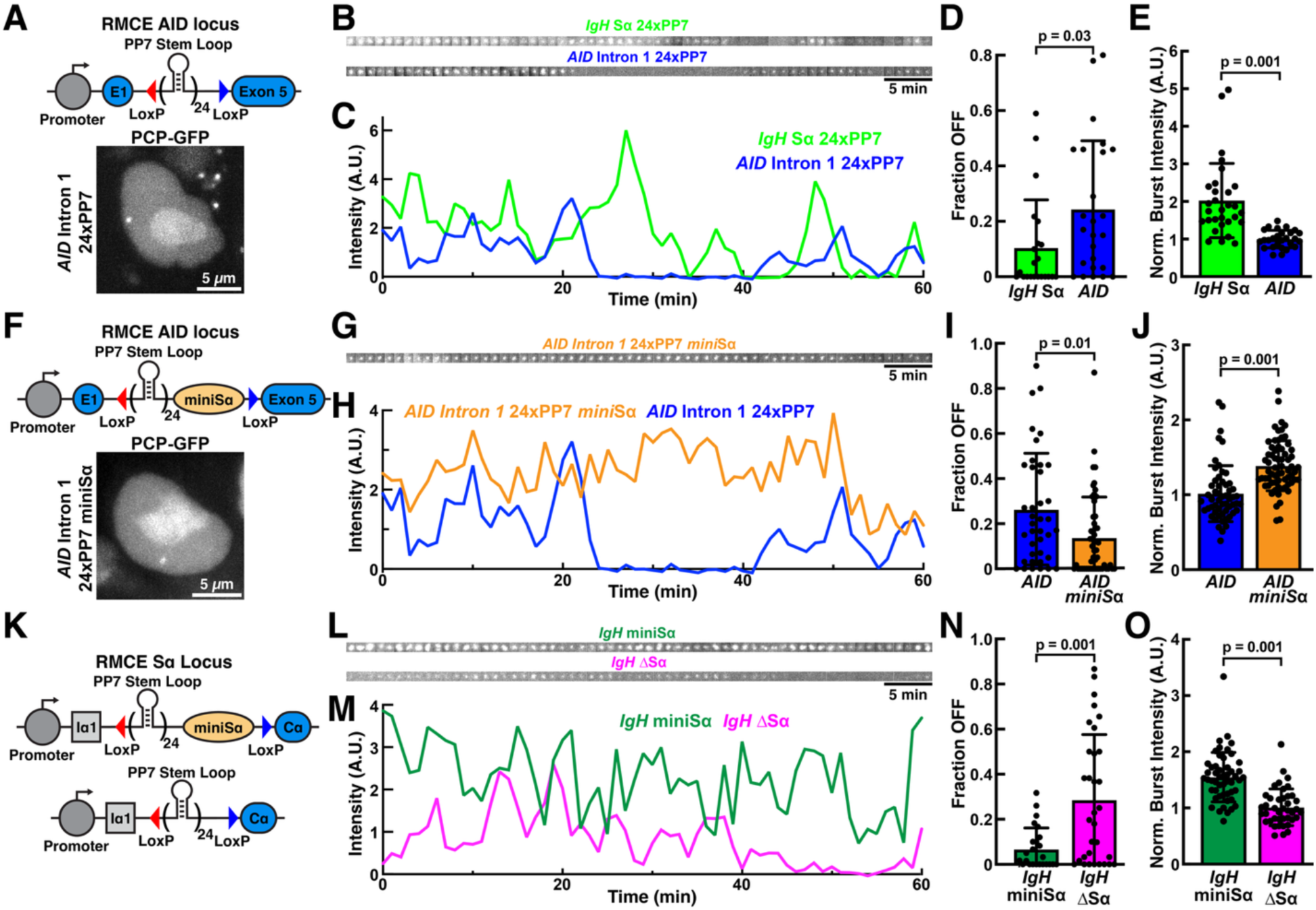
Switch region sequences increase polymerase loading. **(A)** Model of engineered *AID* locus containing a 24xPP7 stem loop array in intron 1 (top). Image of a representative CH12 cell expressing PCP-GFP. **(B-C)** Images and quantification of transcriptional foci intensity of CH12 cells containing a 24xPP7 stem loop array in the *IgH* locus Sα switch region or intron 1 of the *AID* locus. **(D-E)** Quantification of **(D)** the fraction of time without transcriptional signal and **(E)** the peak intensity of transcriptional bursts of traces shown in (C) (N = 22 and 25 cells, Mean±SD, T-Test). **(F)** Model of engineered *AID* locus containing a 24xPP7 stem loop array and the miniSα sequence in intron 1 (top). Image of a representative CH12 cell expressing PCP-GFP. **(G-H)** Images and quantification of transcriptional foci intensity of CH12 cells containing a 24xPP7 stem loop array with and without the miniSα in intron 1 of the *AID* locus. **(I-J)** Quantification of **(I)** the fraction of time without transcriptional signal and **(J)** the peak intensity of transcriptional bursts of traces shown in (H) (N = 39 and 52 cells, Mean±SD, T-Test). **(K)** Model of engineered *IgH* locus containing a 24xPP7 stem loop array with and without the miniSα sequence. **(L-M)** Images and quantification of transcriptional foci intensity of CH12 cells containing a 24xPP7 stem loop array with and without the miniSα sequence in the IgA region of the *IgH* locus. **(N-O)** Quantification of **(N)** the fraction of time without transcriptional signal and **(O)** the peak intensity of transcriptional bursts of traces shown in (M) (N = 25 and 32 cells, Mean±SD, T-Test).

Previous work has demonstrated that the I promoter can be replaced by a heterologous promoter and still support CSR. Therefore, the key functional features of germline transcription may reside in other elements of the transcription unit, such as the switch region. To test the impact of the switch region on transcription, we inserted a minimal Sα switch region (a synthetic fragment containing 14 copies of the consensus 80 nucleotide repeat (Han et al., 2011), hereafter termed miniSα) preceded by the 24xPP7 stem loop array into the first intron of the *AID* locus (Fig. 3F). The presence of the miniSα sequence significantly reduced the transcriptional off-time and increased the burst intensity, relative to the *AID* locus only containing the 24xPP7 stem loop array (Fig. 3G-J, Movie S11), resulting in persistent transcriptional foci similar to those observed at the Sα locus. This demonstrates that the switch region repeats are sufficient to increase polymerase loading, despite being located ∼2 kb downstream of the transcriptional start site. To further confirm this conclusion, we removed the Sα switch region from the *IgH* locus or replaced it with miniSα in either orientation (Fig. 3K). Transcriptional dynamics in the presence of miniSα in the physiological orientation were comparable to the wildtype Sα switch region, while deletion of the switch region sequences resulted in an increase of transcriptional off-time and a reduction in the transcriptional burst intensity (Fig. 3L-O, Movie S12-13). Moreover, inversion of the miniSα sequence in the *AID* or *IgH* locus significantly reduced the intensity of nascent transcript signals (Fig. S3B,C).

In total, these results demonstrate that the Sα switch region sequence in its correct orientation promotes transcriptional activity by increasing polymerase loading, leading to the persistent association of nascent germline transcripts with the *IgH* locus.

### Analysis of the single molecule dynamics of AID in CH12 cells

Our results demonstrate that the switch region sequences are highly transcribed which results in the tethering of multiple nascent germline transcripts to the *IgH* locus and likely exposing single stranded DNA. Since AID has high affinity for switch region RNAs (Qiao et al., 2017; Zheng et al., 2015), the nascent RNA tethered to the switch region could directly recruit AID to the *IgH* locus during CSR.

To analyze the sub-cellular localization and movements of AID in cells with single-molecule sensitivity, we inserted the HaloTag at the C-terminus of the AID coding sequence in the endogenous *AID* locus of CH12 cells (Fig. S4A). Homozygous insertion of the HaloTag was verified by PCR, and exclusive expression of AID-HaloTag (AID-Halo) was confirmed by western blot (Fig. S4B,C). Exposure of CH12 cells to CIT resulted in rapid induction of AID-Halo expression. The AID protein was detected in most cells in 4-6 hours and peaked around 12 hours after stimulation (Fig. S4D-F). To test functionality of the HaloTagged AID, we carried out CSR assays and found that AID-Halo is fully functional, supporting similar levels of CSR as the untagged AID (Fig. S4G,H).

3D imaging of living CH12 cells expressing AID-Halo confirmed that AID is predominantly localized to the cytoplasm (Fig. 4A). We did not observe noticeable localized enrichment of AID at the actively transcribed *IgH* locus (Fig. 4A, Fig. S4I) or any other nuclear region. To investigate the diffusion dynamics of AID, we carried out live cell single-molecule imaging experiments of

**Figure 4.**
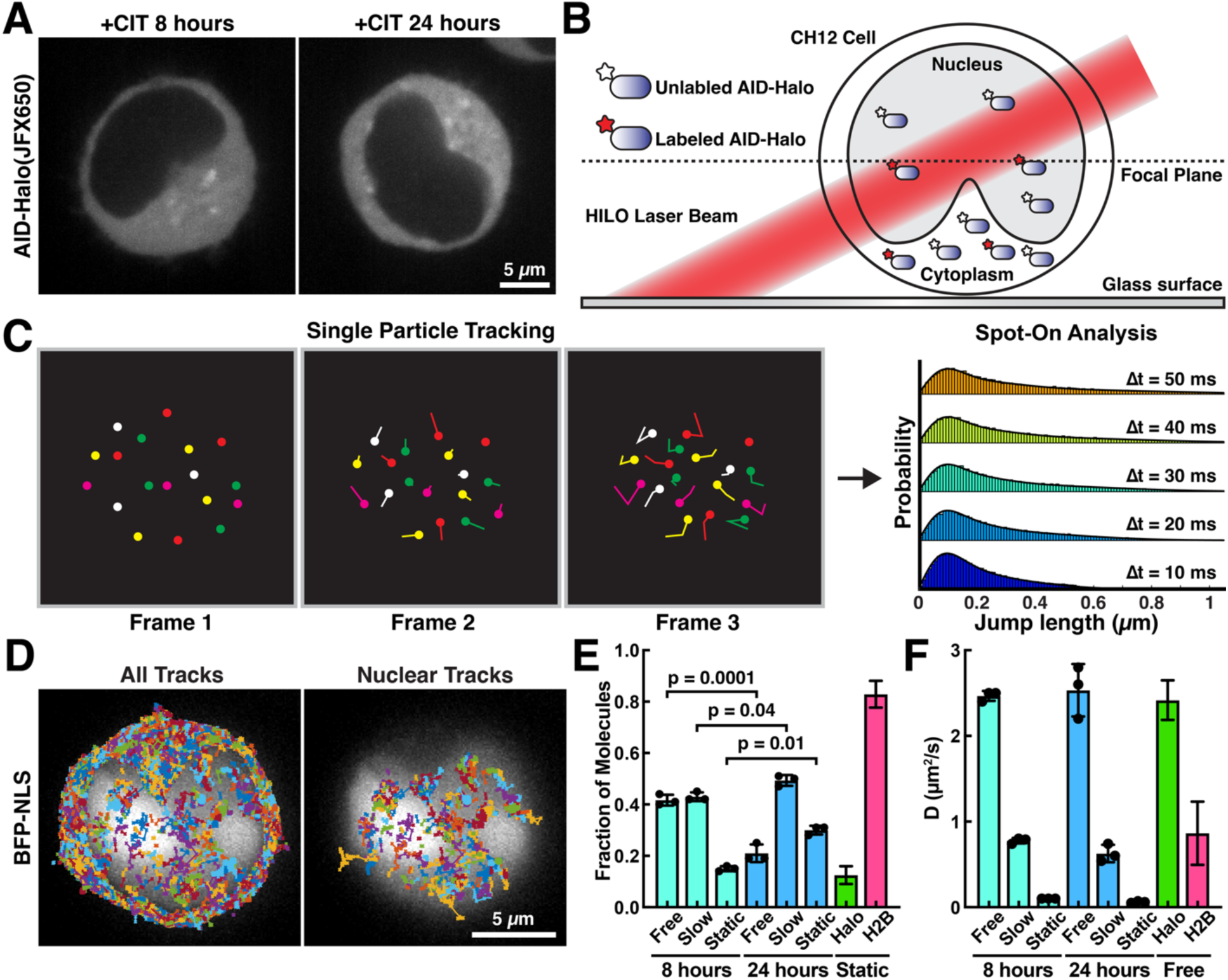
Single-molecule dynamics of nuclear AID-Halo during CSR. **(A)** Images of living CH12 cells expressing AID-Halo from the endogenous AID locus stained with JFX650 HaloTag-ligand. **(B)** Experimental approach for live cell single-molecule imaging in CH12 cells. **(C)** Approach to analyze single particle tracking data using the Spot-On tool. **(D)** Single-molecule trajectories of AID-Halo (JFX650) overlayed on nuclei detected using BFP-NLS. **(E-F)** Spot-On analysis showing the **(E)** fraction of particles and **(F)** diffusion coefficients of molecules in distinct mobility states (Free, Slow, Static) generated from HaloTag and Halo-H2B tracks, or AID-Halo trajectories after CSR induction using CIT for 8 and 24 hours (N = 3 biological replicates, >50 cells per replicate, Mean±SD, One-Way ANOVA).

AID-Halo sparsely labeled with JFX650 and generated trajectories of nuclear AID-Halo molecules using single-particle tracking (Figure 4B-D, Movie S14). As controls for freely diffusing and chromatin bound proteins, we expressed the HaloTag alone or a HaloTag-histone H2B fusion protein in CH12 cells (Movie S15-16). Step size distributions derived from automated single-particle tracking were analyzed using the Spot-On tool (Hansen et al., 2018), which determines the diffusion coefficients and fraction of molecules in distinct molecular states (Fig. 4C). As expected, the HaloTag protein diffused rapidly through the nucleus (D_free_ = 2.4 µm^2^/s) and only a small fraction of molecules was static (F_bound_ = 12%, Fig. 4E-F, Movie S15). In contrast, the majority of Halo-H2B molecules was static (F_bound_ = 83%, Fig. 4E, Movie S16), consistent with its association with chromatin. Analysis of the diffusion dynamics of nuclear Halo-AID 8 hours after CSR induction revealed that the step size distributions of AID movements fit well to a three-state model composed of rapidly (D_free_ = 2.5 µm^2^/s) and slowly diffusing (D_slow_ = 0.77 µm^2^/s) as well as static (D_static_ = 0.1 µm^2^/s) AID molecules (Fig. 4F, Fig. S4J, Movie S14). Rapidly diffusing signals likely represent individual AID molecules exploring the nucleoplasm by Brownian motion, while static particles represent AID molecules associated with chromatin, including the *IgH* locus. Slowly diffusing AID signals could report on AID molecules that are part of large macromolecular complexes, for example AID bound to nuclear RNA molecules. It is important to note that the diffusion co-efficient of slowly diffusion AID (D_slow_ = 0.63 µm^2^/s) closely corresponds to that of mobile germline transcripts at 24 hours (D_free_ = 0.76 µm^2^/s, Fig. 1M), which is consistent with AID binding to these RNA molecules. Eight hours after induction of CSR, AID-Halo was largely mobile (F_free_ = 42%, F_slow_ = 43%) with a small fraction of AID molecules being static (F_static_ = 15%) (Fig. 4E). Importantly, the fraction of static AID-Halo molecules at this time point is only slightly higher than that of the HaloTag alone. We therefore conclude that, 8 hours after CSR induction, AID is not specifically associated with chromatin. After stimulation of CSR for 24 hours, the fraction of static AID-Halo molecules increased significantly to F_bound_ = 30%, the fraction of highly mobile AID-Halo molecules was significantly decreased to F_free_ = 21%, while the fraction of slowly diffusing AID-Halo particles significantly increased (F_slow_ = 49%) (Fig. 4E). These observations are consistent with the recruitment of AID to chromatin 24 hours after CSR induction.

Together, these results demonstrate that the limited number of nuclear AID molecules exist in three states (freely diffusing, slowly diffusing, and chromatin bound), and that the fraction of chromatin-bound AID is significantly increased over the course of CSR, which could in part reflect the recruitment of AID to the *IgH* locus.

### AID recruitment to chromatin requires zinc coordination in the AID active site

To further dissect the molecular mechanism underlying AID recruitment to chromatin, wildtype and mutant variants (H56A, E58A, and R190X) of AID fused to the HaloTag were introduced into one allele of the endogenous *AID* locus via RMCE in an engineered CH12 cell line in which the second *AID* allele has been knocked out (Fig. 5A,B, Fig. S5A,B). This approach supported robust AID expression and class switch recombination (Fig. S5C-E). H56 and E58 are critical residues in the active site of AID involved in coordination of a zinc ion (H56) and acid base-mediated catalysis (E58), respectively (Qiao et al., 2017) (Fig. 5A). The R190X variant of AID, caused by a premature stop codon that eliminates the nuclear export signal at the C-terminus of AID, was found in patients suffering from Hyper-IgM syndrome with a dominant negative phenotype (Imai et al., 2005). Consistent with their importance for AID function, the H56A, E58A, and R190X mutations eliminated CSR (Fig. S5F). The overall nuclear distribution of H56A and E58A AID variants was similar to that of wildtype AID, with AID being evenly distributed throughout the nucleus without detectable sites of accumulation (Fig. 5C). Nuclear abundance of R190X AID was increased, consistent with the mutation affecting the nuclear export signal in AID (Geisberger et al., 2009). The R190X was depleted from large circular areas in the nucleus, which likely correspond to nucleoli (Fig. 5C). Analysis of the nuclear diffusion dynamics of the AID mutants revealed that E58A and R190X behave similarly to wildtype AID, with an increase in the chromatin bound fraction and reduction of the fraction of freely diffusing AID molecules at 24 hours compared to 8 hours of CSR induction (Fig. 5D-E, Movies S17-19). In contrast, the diffusion dynamics of H56A AID were unchanged at the 24-hour time point and comparable to wildtype AID at 8 hours after CSR induction (Fig. 5D-E). These observations demonstrate that the chromatin recruitment of AID is reduced by the H56A mutation and suggest that zinc ion coordination could potentially contribute to substrate binding and the immobilization of AID.

**Figure 5.**
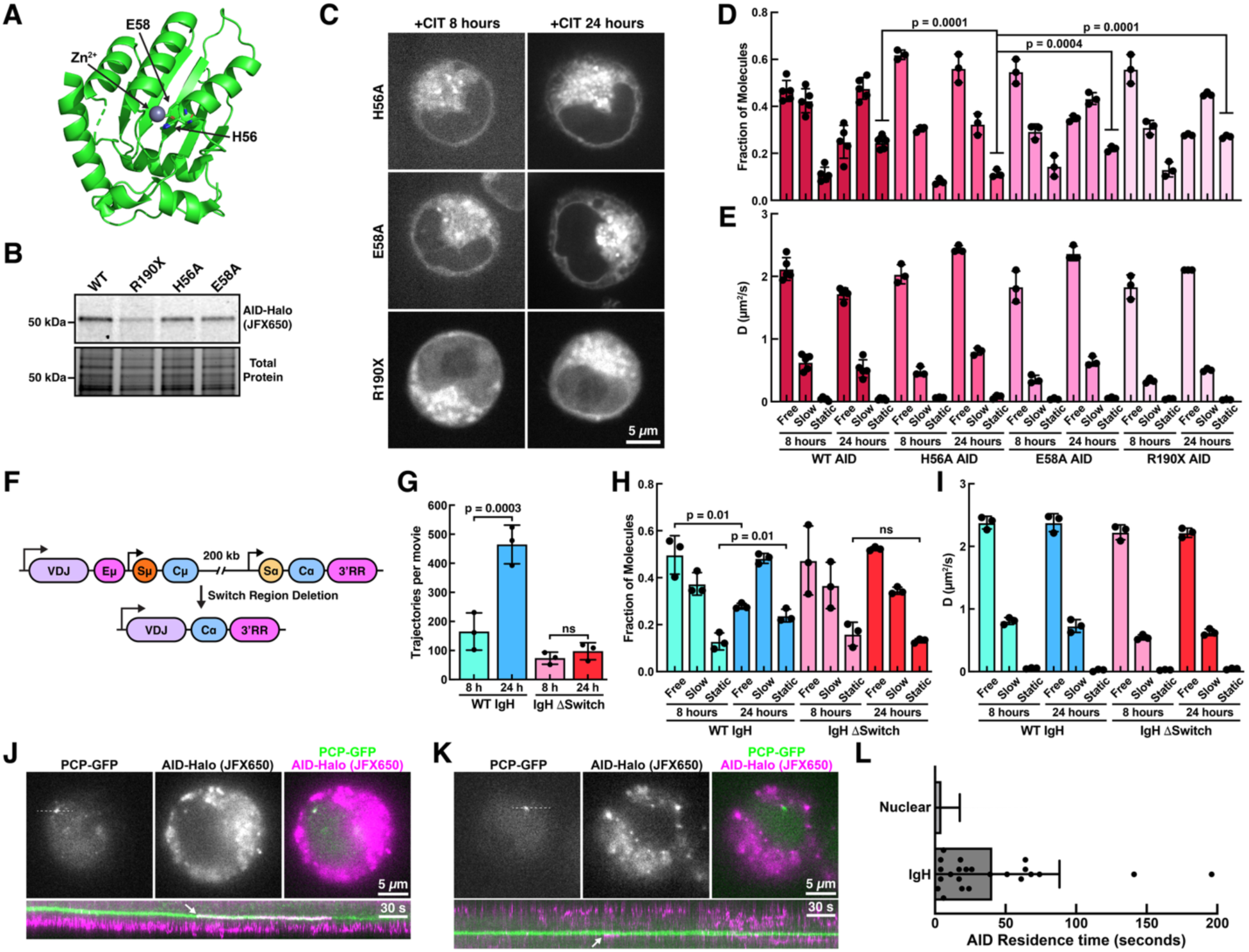
Molecular determinants of AID recruitment to chromatin and the *IgH* locus. **(A)** Structure of AID (adapted from PDB:5W0Z) indicating the location of H56 and E58. **(B)** Fluorescent gel (top) and total protein stain (bottom) of SDS-PAGE gel loaded with cell lysates generated from CH12 cells expressing HaloTagged wildtype, R190X, H56A, or E58A AID labeled with JFX650 HaloTag-ligand. **(C)** Images of living CH12 cells expressing AID-Halo variants from the endogenous AID locus stained with JFX650 HaloTag-ligand. **(D-E)** Spot-On analysis showing the **(C)** fraction of particles and **(D)** diffusion coefficients of molecules in distinct mobility states (Free, Slow, Static) generated from AID-Halo trajectories after CSR induction using CIT for 8 and 24 hours from CH12 cells expressing HaloTagged wildtype, R190X, H56A, or E58A AID (N = 3-5 biological replicates, >60 cells per replicated, Mean ± SD, one-way ANOVA). **(F)** Genome editing strategy to delete the switch regions from the *IgH* locus. **(G)** Quantification of the number of nuclear AID-Halo trajectories in control CH12 cells and CH12 cells lacking the switch region sequences after induction of CSR using CIT for 8 and 24 hours. **(H-I)** Spot-On analysis showing the **(H)** fraction of particles and **(I)** diffusion coefficients of molecules in distinct mobility states (Free, Slow, Static) generated from AID-Halo trajectories after CSR induction using CIT for 8 and 24 hours from control CH12 cells and CH12 cells lacking the switch region sequences (N = 3 biological replicates, >60 cells per replicated, Mean ± SD, T-Test). **(J-K)** Single frames of time-lapse movies acquired at 1 frame per second of living CH12 cells with a 24xPP7 stem loop array integrated into the Sα switch region expressing PCP-GFP and AID-Halo labeled with JFX650 (top). Kymographs along the white dashed line over 400 frames of the time lapse movies (bottom). **(L)** Quantification of the residence time of AID-Halo at the IgH locus and all nuclear locations.

### AID recruitment to chromatin requires switch regions

To determine whether the switch regions of the *IgH* locus and potentially their RNA products contribute to the chromatin recruitment of AID, we deleted the genomic region containing all switch regions from both *IgH* alleles in CH12 cells expressing AID-Halo from the endogenous *AID* locus (Fig. 5F, Fig. S5G,H). Single-particle tracking demonstrated that the number of nuclear AID trajectories in control cells with an intact *IgH* locus increased over the course of CSR (Fig. 5G). In contrast, in the cells lacking any switch region sequence, the number of nuclear AID trajectories was not increased after 24 hours of CSR induction relative to the 8-hour time point (Fig. 5G). Consistent with the results described above (Fig. 5E), in control cells the fraction of static AID molecules was increased 24 hours after CSR induction compared to 8 hours after CIT stimulation (Fig. 5H-I). However, the diffusion properties of AID-Halo in cells lacking all switch regions were identical at 8 and 24 hours after CSR induction, with a low fraction of chromatin bound AID molecules (F_static 8 hours_ = 16%, F_static 24 hours_ = 13%) and a large fraction of freely diffusing AID particles (F_free 8 hours_ = 47%, F_free 24 hours_ = 52%) (Fig. 5H-I, Movie S20). These results demonstrate that switch region transcription is required for the changes in AID dynamics during CSR, which is consistent with AID interacting with germline transcripts at the *IgH* locus or elsewhere in the nucleus.

### AID specifically associates with actively transcribed *IgH* locus

To analyze the specific recruitment of AID to the *IgH* locus, we carried out multi-modal single-molecule imaging, simultaneously visualizing the *IgH* locus marked by nascent 24xPP7 labeled Sα transcripts and AID-Halo molecules (Fig. 5J-K, Movie S21). Capturing images at one-second intervals over ten minutes allowed us to detect rare but long-lasting interactions between AID and the *IgH* locus (Fig. 5J-K, Movie S21). These interactions were defined as co-localizations of PCP-GFP and AID-Halo signals. In all cases, the co-localized signals co-migrated in the nucleus for multiple imaging frames (Fig. 5J-K, Movie S21), raising our confidence that they report on bona fide AID interactions with the *IgH* locus. AID molecules remained bound to the *IgH* locus for an average of 40 seconds, a duration nearly an order of magnitude longer than the residence time of static AID molecules elsewhere in the nucleus, which lasted on average 4.7 seconds. AID interactions with the *IgH* locus were exceedingly rare. We imaged 288 cells for a total of 118,799 seconds and AID localizations to the *IgH* locus spanned a total of 938 seconds, corresponding to an occupancy of 0.7% of the time. However, because we only label a small fraction of AID-Halo molecules in our single-molecule imaging experiments and considering the fraction of labeled AID-Halo molecules (4±1%, Fig. S5I), we conclude that the *IgH* locus is occupied by AID approximately 18% of the time during CSR. This prolonged binding demonstrates that AID has a higher affinity for the *IgH* locus compared to other random genomic locations.

## DISCUSSION

Our real-time analysis of germline transcription revealed that the Sα switch region enhances polymerase loading, even though it is located 2kb downstream of the transcriptional start site. In addition, our results demonstrate that transcription rapidly proceeds through the repetitive switch region, which contradicts previous models that proposed slow or even stalled transcription within the switch region (Rajagopal et al., 2009; Wang et al., 2009). We currently do not know how switch region promotes polymerase loading. However, since its correct orientation is critical for this effect, it is possible that transient R-loop formation, or effects on DNA topology, for instance changes in DNA supercoiling, induced by the switch region or its transcription could alter the upstream promotor structure to increase the number of RNA polymerases loaded during transcriptional bursting. Importantly, our observations suggest that R-loops at switch regions are either transient or do not hinder transcriptional elongation. Several transcription factors including Spt5 (Maul et al., 2014; Pavri et al., 2010), PAF1 (Willmann et al., 2012), and ELOF1 (Dai et al., 2025; Wu et al., 2025) were found to be important for efficient CSR. These factors associate with paused RNA polymerases at promoter proximal regions and regulate their transition into active transcriptional elongation. It is therefore possible that Spt5, PAF1, and ELOF1 control the burst frequency and amplitude of germline transcription to promote optimal CSR.

The high levels of germline transcription could contribute in two distinct ways to AID mediated deamination of switch regions. First, rapid transcription of the switch region by densely loaded polymerases may facilitate the exposure of single strand DNA to AID, potentially by preventing re-annealing of the DNA duplex after a polymerase has passed (Fig. 6). Even small changes in the number of polymerases loaded or the rate of transcription could limit the number of hotspots available for AID to target, which would reduce the efficiency of CSR. Secondly, high levels of transcription lead to the formation of a dynamic RNA hub seeded by multiple nascent germline transcripts associated with the *IgH* locus (Fig. 6). Our RNA imaging demonstrates that on average 5 but up to 12 Sα switch region containing transcripts simultaneously associate with the *IgH* locus. Together with RNAs derived from Sµ, which are in proximity to Sα due to the 3D chromatin structure at the *IgH* locus (Wuerffel et al., 2007; Zhang et al., 2019), the total number of germline transcripts within the RNA hub likely ranges from 10-20 molecules. In addition, we observed mobile, intronic RNAs containing the switch region sequences that can potentially rebind the RNA hub, suggesting that it may form a dynamic condensate allowing association and dissociation of RNA molecules, which may facilitate rebinding of RNA-bound AID to the *IgH* locus.

**Figure 6.**
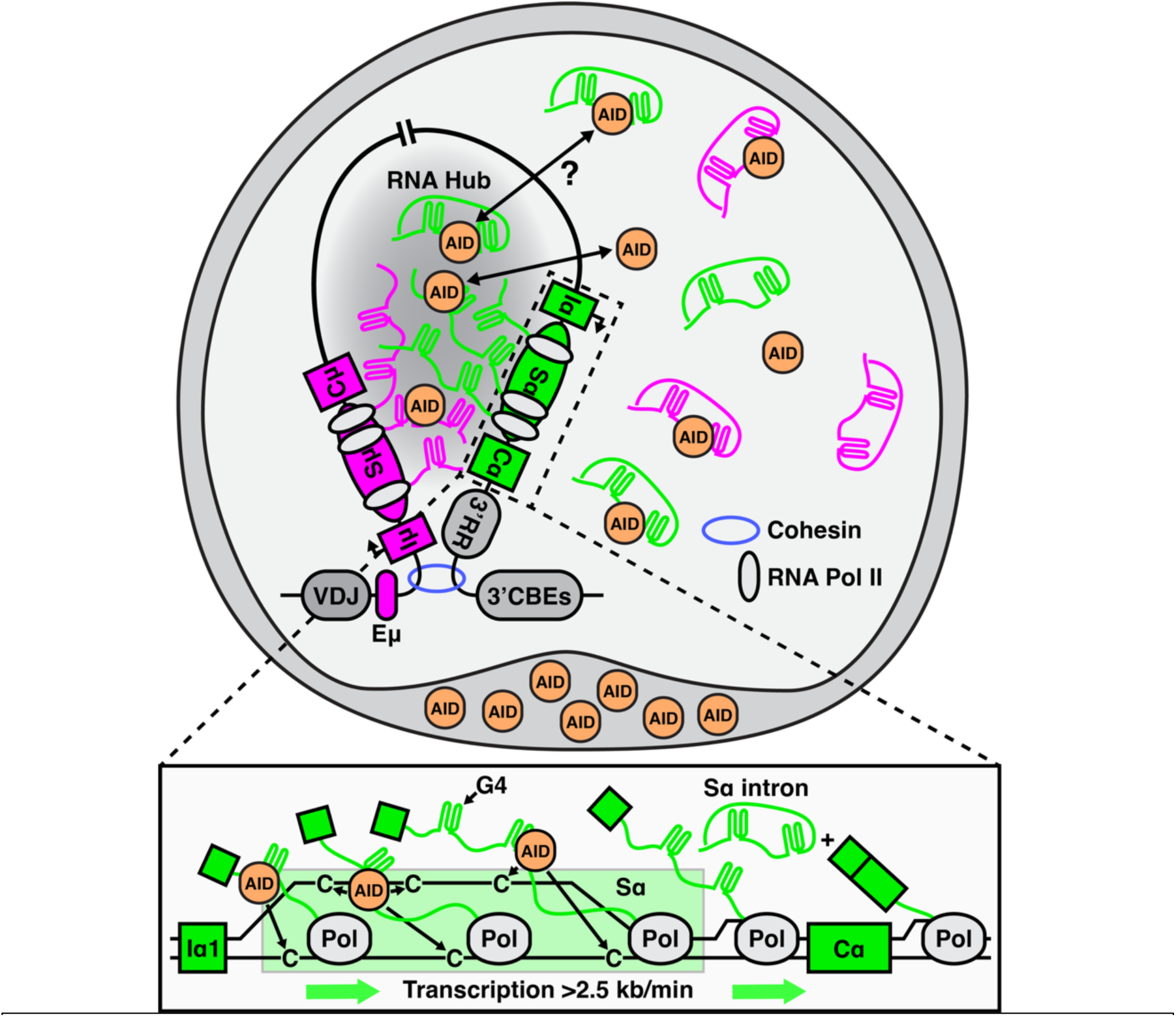
Model for RNA mediated recruitment of AID to the *IgH* locus. Rapid transcription of the switch region by dense bursts of RNA polymerases exposes single-stranded DNA and result in a large number of nascent germline transcripts directly tethered to the *IgH* locus. AID directly associates with the germline transcripts to facilitates its recruitment to the *IgH* locus and allow it to target the single-stranded switch region DNA exposed by high transcriptional activity.

We propose that the RNA hub forms a landing platform to recruit AID to the *IgH* locus (Fig. 6). Our results demonstrate that the dynamics of nuclear AID movement are altered over the course of CSR. Importantly, these changes coincide with an increase in the amount nuclear germline transcripts that can be totally eliminated simply by deleting all switch regions in the cell. These observations strongly suggest that AID directly associates with germline transcripts *in vivo*. This interpretation is further supported by the close correspondence of the diffusion coefficient of mobile RNA molecules and slowly moving nuclear AID particles. Previous work by others has shown that AID has a high affinity for G4 containing RNA and branched nucleic acid structures, which can be formed by switch region sequences (Qiao et al., 2017; Zheng et al., 2015). Therefore, the RNA hubs likely contain a large number of AID binding sites capable of recruiting AID to the *IgH* locus. Once AID localizes to the *IgH* locus, it is retained for an extended period (on average 40 seconds) potentially allowing deamination of multiple cytosine bases within the switch regions, while binding to the nascent germline transcript (Liu et al., 2021).

Our study suggests that the switch region sequence has three critical features to support RNA hub formation for AID recruitment and subsequent cytosine deamination. It contains AID hotspots, it promotes high levels of transcription, and its RNA product forms secondary structures that bind to AID. These properties might not be unique to GC rich switch regions with high G-content on the non-template strand, such as switch regions in mice and humans. For example, the A-T rich switch region from *Xenopus Laevis* (Zarrin et al., 2004) or an inverted Sµ fragment from mice (Dong et al., 2015) could have similar properties which partially support CSR. These sequences also contain many AID hotspot motifs that may attract AID. It is tempting to speculate that the RNA hub represents transcriptional condensates containing both acceptor and donor switch region transcripts, which could contribute to juxtaposition of distant switch regions to facilitate the deletional DNA break joining required for productive CSR (Fig. 6) (Dong et al., 2015). This model aligns with the cohesion-mediated loop extrusion in CSR, which brings distant switch regions into close proximity (Zhang et al., 2019). Loop extrusion was proposed to be impeded at transcribed switch regions to bring together donor and acceptor switch regions (Zhang et al., 2019). In this sense, the RNA hub could constitute a structural and biochemical microenvironment of a “recombination center” (Yu, 2021; Zhang et al., 2021) to create a dynamic nuclear compartment where multiple layers of regulation converge to choreograph AID activity and DSB formation to facilitate orientation-specific DNA end joining during CSR. A similar mechanism involving condensate formation and loop extrusion has been proposed for establishing enhancer promotor contacts in the context of transcriptional regulation (Sharp et al., 2022). Therefore, our findings might extend beyond antibody diversification and have wide ranging implications in transcriptional regulation and 3D genome organization.

## MATERIALS AND METHODS

### Oligonucleotides

Oligonucleotides were purchased from Millipore-Sigma and Integrated DNA Technologies (IDT). Synthetic guide RNAs were purchased from IDT.

List of sgRNAs (5’ target–PAM 3’)

KY1139 (5’ GGATTTTGAAAGCAACCTCC–TGG 3’), C-terminus of AID.

KY1284 (5’ AAGGACAGTGCTTAGATCCG–AGG 3’), 5’ of Sµ.

KY1156 (5’ CTCTGCTCATTGGTACACTG–AGG 3’), 5’ of Sα.

KY1268 (5’ TGGAGCGCTAGACTGCTCAG–GGG 3’), 3’ of Sα.

KY1278 (5’ CAACTACCCTTTTGAGACCG–AGG 3’), 5’ of Eµ.

### CH12 cell culture and class switch assay

CH12 cells were cultured in RPMI 1640 medium supplemented with 10% (vol/vol) fetal bovine serum and 50 µM β-mercaptoethanol. To induce CSR, CH12F3 cells were seeded at 5 x10^4^ cells/ml in culture medium containing 0.5 µg/ml anti-CD40 antibody (Ab) (clone HM40-3; eBioscience), 5 ng/ml IL-4 (R&D Systems), and 0.5 ng/ml TGF-β1 (R&D Systems) and grown for 72 hr unless otherwise specified. Cells were stained with a CF633 (Biotium)-conjugated rat anti-mouse IgA monoclonal antibody (556969; BD Biosciences) and analyzed on an Attune CytPix flow cytometer. Flow cytometry data were analyzed using the Flowjo software (BD Biosciences). CSR efficiency was determined as the percentage of IgA-positive cells.

### CRISPR/Cas9 genome edition

Deletion of the *IgH* non-productive allele was performed as previously described (Kim et al., 2016) as the first step for generating all cell lines in this study. **For tagging AID at its C-terminus**, a linear fragment containing the Halo-3xFLAG coding region flanked by two homology arms (290 bp and 1603 bp) was used as the homology-directed repair template (HDRT) (Fig. S3A). Ribonucleotide protein complex (RNP) was assembled by incubating the purified Cas9 protein and a synthetic guide RNA (KY1139). About 500000 CH12 cells were transfected (1450V, 10ms, 3 pulses) with 50 pmoles of Cas9RNP and 200 ng of HDRT in a 10µl tip using a Neon electroporation device (ThermoFisher Scientific) according to the manufacturer’s instructions. After a 24hr recovery, cells were subcloned by limited dilution in a 96-well plate. Single clones were picked after 7 days. A replicate set of clones were used to test for Halo expression after 24hr CIT stimulation followed by labeling with the JF635 halo ligand (Janelia Research Campus). Halo positive clones were genotyped by PCR (Fig. S3A,B). Homozygous clones were further confirmed by western blotting (anti-AID) for the exclusive expression of the AID-halo fusion protein after 24hr CIT stimulation (Fig. S3C). **For tagging germline transcripts**, Cas9RNP and HDRT was transfected to CH12 or its derivatives by Neon transfection. To insert the 24xPP7 array upstream of Sα, Cas9RNP (KY1156) and an HDRT containing a 1311 bp left homology arm and a 980 bp right homology arm flanking an array of 24 PP7 stem loops (1.5 kb XhoI fragment from Addgene plasmid # 61762) was co-transfected to WT or AID-Halo CH12 cells (Fig. S1A,B). To insert the 24xMS2 array upstream of Sµ, Cas9RNP (KY1284) and an HDRT containing a 852 bp left homology arm and a 467 bp right homology arm flanking an array of 24 MS2 stem loops (1.3 kb Bgl II to Sac I fragment from Addgene plasmid # 61762) was transfected to cells already had Halo inserted at AID and 24xPP7 inserted upstream of Sα (Fig. S1E). To insert the 24xMS2 downstream of Sα, Cas9RNP (KY1268) and an HDRT containing a 743 bp left homology arm and a 493 bp right homology arm flanking an array of 24 MS2 stem loops (1.3 kb Bgl II to Sac I fragment from Addgene plasmid # 61762) was transfected to cells that had Halo inserted at the C-terminus of AID and 24xPP7 inserted upstream of Sα (Fig. S2A). In each case, transfected cells were single cloned and screened by PCR to identify clones with the correct integration. To generate cells with no switch region, AID-Halo cells were transfected of Cas9RNPs assembled with KY1268 and KY1278 to delete all switch and constant regions on the productive allele (Fig. S4G). Transfected cells were single cloned and screened by PCR to identify clones with the correct deletion.

### RMCE at AID locus

Homology mediated gene targeting was performed in our previously established AID knockout cell line (Ramachandran et al., 2016) to insert an RMCE cassette replacing most of the AID coding sequences (exons 2-4) (Fig. S4A-C). The resulting cell line contains two null alleles, one of which contains an RMCE cassette. The RMCE cassette is made of a positive-negative selection marker (PGK-Puro-miniTK) flanked by a pair of heterologous LoxP sites. Co-transfection of a Cre-recombinase expression vector and an exchange plasmid (carrying test sequences flanked by the same pair of heterologous LoxP sites) results in swapping of cassettes between the chromosome and the exchange plasmid. Successful RMCE events were counter-selected with gangcyclovir. In this design, AID variants carrying mutations to any but the first three amino acid residues (encoded by Exon 1) can be monoallelicly expressed from the endogenous *AID* locus (Fig. S4A-F). Similarly, switch sequences can be targeted into the *AID* locus (intron 1) to test their impacts on transcription elongation or stalling (Fig. 4C).

### RMCE at the *IgH* Sα locus

RMCE at the *IgH* Sα locus was performed in a previously established CH12 cell line as described (Han et al., 2011).

### Measurement and quantification of in-gel fluorescence

80,000 CH12F3 cells were stimulated with CIT for 0-24 hours. Cells labeled with 100 nM JFX650 HaloTag-ligand for 30 minutes in culture media, spun down at 1000 PPM for 5 minutes, media with dye was remove, and cells were washed with PBS. Next, cells were spun down and lysed in 100 μL Laemmli sample buffer (BioRad) and boiled for 5 min at 95°C. To measure the fraction of AID molecules that we are labeled during single-molecule imaging experiments, we used AID single-molecule imaging labeling conditions (0.6 nM JFX650 HaloTag-ligand for 30 seconds). Samples were loaded and separated on Mini-PROTEAN TGX stain-free gels (BioRad) and fluorescence and then total protein abundance were measured using a BioRad Chemidoc, using the Cy5.5 filter and stain-free gel detection after 45 seconds of UV activation, respectively. Protein loading was normalized using stain-free total protein detection. This approach allowed us to ensure accurate quantification and comparison of protein expression levels. All gels were analyzed using ImageQuant TL 8.2. For quantification of the labeling efficiency in single-molecule imaging experiments, the fluorescence signal was normalized to the total protein signal to normalize for differences in loading and cell number. All experiments were performed in triplicate.

### Immunoblotting and Antibody

Cell lysates were prepared in RIPA buffer (50 mM Tris•HCl, pH 7.4, 150 mM NaCl, 1% Triton X-100, 0.5% sodium deoxylcholate, 0.1 % SDS and 1 mM EDTA) and resolved by sodium dodecyl sulfate polyacrylamide gel electrophoresis. Proteins were transferred to a 0.2µm PVDF membrane and probed with a Rat anti-AID antibody (14-5959-82, eBioscience).

### Live-Cell Single-Molecule Imaging (L-C S-M) to track AID dynamics

For live-cell single-molecule imaging, CH12F3 cells were transfected with a nuclear-localized blue fluorescent protein (BFP-NLS) to delineate the nuclear boundary. Cells were then stimulated with CIT for 8 and 24 hours to induce CSR and AID expression. On the day of imaging, cells were sparsely labeled with 2 nM of JFX650 dye for 1 minute, were span down at 1000 RPM for 3 minutes in falcon tubes and washed three times with media, last wash included an incubation for an additional 15 min at 37 °C (5% CO2) in CIT media to remove excess dye. Prior to imaging, 250000 cells were transferred into 24-well glass bottom dishes (Cellvis, P2 4-1.5H-N) and span down at 1000 RPM for 5 minutes to prevent B cells from excessive movements during imaging. Movie acquisition was performed in a cellVivo humidified incubation chamber at 37° C supplied with 5% CO2. Before the acquisition, cells were allowed to equilibrate to 37° C for about 15-30 minutes. Acquisition was performed on Olympus IX83 inverted microscope equipped with a 4-line cellTIRF illuminator (405 nm, 488 nm, 561 nm, 640 nm lasers), an Excelitas X-Cite TURBO LED light source, an Olympus UAPO 100x TIRF objective (1.49 NA), an Olympus UPlanApo 60x TIRF oil-immersion objective (1.50 NA), a CAIRN TwinCam beamsplitter, two Andor iXon Ultra EMCCD cameras or two Hamamatsu Orca BT Fusion cameras, or a Hamamatsu Orca QUEST (2x2 binning) camera. For each movie we recorded 1500 frames at 146 or 172 frames per second. All experiments were carried out in triplicate or more with at least 60 cells analyzed for each experiment.

### Single-particle Tracking Analysis

Single-particle tracking was carried out using MATLAB R2023a using a batch parallel-processing implementation of SLIMfast) using the following settings: Exposure Time = 1/frame rate, NA = 1.49, Pixel Size = 0.1083 µm (BT Fusion) or 0.1533 µm (QUEST), Emission Wavelength = 664 nm, D_max_ = 5 µm^2^/s, Number of gaps allowed = 2, Localization Error = 10^-5^, Deflation Loops = 0 ^28^. Diffusion coefficients and state distributions were determined using the Spot-On tool using the following settings: TimeGap = 1/frame rate, dZ = 0.700 µm, GapsAllowed = 2, TimePoints = 8, JumpsToConsider = 4, BinWidth = 0.01 µm, PDF-fitting, D_Free1_3State = [1 25], D_Free2_3State = [0.1 1], D_Bound_3State = [0.0001 0.1] (Hansen et al., 2018). All experiments were carried out in triplicate or more and at least 60 cells were analyzed per replicate. Track information from individual cells was pooled for Spot-On analysis.

### AID Residence Time Analysis

To determine the residence time of AID at the *IgH* locus images of Halo-AID labeled with JFX650 and the IgH locus marked by the nascent Sα switch region transcript where simultaneously imaged at a rate of 1 frame per second. The residence time of static nuclear AID molecules was determined by single-particle tracking. To limit the analysis to static molecules the diffusion coefficient during the tracking step was constrained to D < 0.01 µm^2^/s and maxOffTime = 2 frames, as previously described (Schmidt et al., 2016). Using this approach the residence time is equal to the track length. Due to their rare occurrence *AID* molecules at the *IgH* locus were analyzed by manual inspection to avoid false positives.

### Real-time Transcription Imaging

500,000 CH12F3 cells carrying 24xPP7 and/or 24xMS2 stem loop arrays were transfected with plasmids encoding PCP-sfGFP and/or MCP-mScarlet (0.5 µg total DNA, at a ratio of 4:1 when both PCP-sfGFP and MCP-mScarlet are included). Approximately 8 hours after electroporation, cells were spun down (180xg for 3 minutes) and resuspended in 2 mL of CIT-containing medium. At the time of imaging (8-24 hours post CIT induction), 50,000–100,000 cells were transferred into glass-bottom dishes (Cellvis, P24-1.5H-N) and centrifuged at 180xg for 5 minutes to ensure that the suspension B cells settled at the bottom of the plate and could be imaged. Once on the microscope, the sample was warmed and maintained at 37 °C with 5% CO_2_ and 95% humidity for the duration of the experiment. Imaging began at least 30 minutes after the cells were positioned in the microscope-attached incubator to minimize thermal drift. Images were acquired using 3i spinning-disc (Yokogawa) microscope with a SoRa disk, equipped and incubation chamber (temperature, humidity and CO_2_ controlled), four laser lines (445, 488, 561 and 638 nm), a Zeiss C PlanApo 63x/1.42 NA objective, and a Hamamatsu Orca-Fusion BT sCMOS camera. Cells were excited with 488 nm and 561 nm laser lines. A Z-stack of 16-20 planes with 0.5 μm spacing was collected using a 200 ms exposure time for both spectral channels, and data were acquired at a frame rate of 1 Z-stack every 60 seconds for a total of 1 hour.

### Real-time Transcription Analysis

To analyze the intensity of transcriptional foci over time Z-stacks were maximum intensity projected and we determined the maximal pixel intensity of the foci formed by PCP-GFP or MCP-mScarlet for every frame of the movie. Background subtraction was carried out by determining the nuclear PCP-GFP or MCP-mScarlet signal intensity of the first and last frame of each cell analyzed and linearly extrapolating between the two time points to account for photobleaching of the background signal. To normalize the signal intensities between independent experimental replicates all data points of individual replicates were divided by the mean intensity of the experimental group with the lower average signal of the experimental replicate. Transcriptional bursts were defined as peaks in the transcriptional intensity trace that were preceded by a signal reduction of at least 50% from the previous peak value and at least a 2-fold increase of intensity relative to the previous minimum in the intensity trace. The rate of transcription across the Sα switch region was calculated by determining the time points at which the PCP-GFP and MCP-mScarlet traces peaked. If the peaks occurred at the same time the transcriptional burst must have traveled through the 24xPP7 stem loop array, the Sα switch region, and the 24xMS2 stem loop array (4.7 kb) in less than a minute and therefore the rate of transcription would be >5 kb per minute. For PCP-GFP and MCP-mScarlet separated by at least one imaging time point (1 minute), we divided the number of nucleotides incorporated (4.7 kb) by the time difference between the peaks.

## ACKNOWLEDGEMENTS

We thank Dr. Bin Wu for the tdMCP-mScarlet expression vectors, and Dr. Fred Alt for comments and discussion on the manuscript. This work was supported by a NIH grants (R01 AI139039) to K.Y. and (DP2 GM142307) to J. C. S.

## AUTHOR CONTRIBUTIONS

Conceptualization: M.M., J.C.S., and K.Y.; Experiments: M.M., M.K., L.H, J.C.S., and K.Y.; Data Analysis: M.M., M.K., J.C.S., and K.Y.; Writing: Original Draft: M.M., J.C.S., and K.Y.; Writing: Review and Editing: M.M., J.C.S., and K.Y.

## SUPPLEMENTAL MOVIE LEGENDS

**Movie S1.** Live cell imaging movie of CH12 cells containing a 24xPP7 stem loop array upstream of the endogenous Sα switch region expressing PCP-GFP after 8 hours of stimulation with CIT. A single focal plane was acquired with a SoRa spinning disc confocal using a 63x objective, the 2.8x magnification lens, and the Hamamatsu ORCA Quest camera at 2x2 binning at 5 frames per second.

**Movie S2.** Live cell imaging movie of CH12 cells containing a 24xPP7 stem loop array upstream of the endogenous Sα switch region expressing PCP-GFP after 24 hours of stimulation with CIT. A single focal plane was acquired with a SoRa spinning disc confocal using a 63x objective, the 2.8x magnification lens, and the Hamamatsu ORCA Quest camera at 2x2 binning at 5 frames per second.

**Movie S3.** Live cell imaging movie of CH12 cells containing a 24xPP7 stem loop array upstream of the endogenous Sα switch region and a 24xMS2 stem loop array upstream of the Sµ switch region expressing PCP-GFP (left panel, green in merge) and MCP-mScarlet (middle panel, magenta in merge) after 8 hours of stimulation with CIT. A Z-stack of the cells was acquired every minute using a SoRa spinning disc confocal with a 63x objective and a Hamamatsu ORCA-Fusion BT camera without binning. The movie is a maximum intensity projection of the Z-stacks.

**Movie S4.** Live cell imaging movie of CH12 cells containing a 24xPP7 stem loop array upstream of the endogenous Sα switch region and a 24xMS2 stem loop array upstream of the Sµ switch region expressing MCP-mScarlet after 8 hours of stimulation with CIT. A single focal plane was acquired was acquired at 4.5 frames per second using a SoRa spinning disc confocal with a 63x objective and a Hamamatsu ORCA-Fusion BT camera without binning.

**Movie S5.** Live cell imaging movie of CH12 cells containing a 24xPP7 stem loop array upstream of the endogenous Sα switch region expressing PCP-GFP without an NLS after 8 hours of stimulation with CIT. A single focal plane was acquired at 10 frames per second using a SoRa spinning disc confocal with a 63x objective and a Hamamatsu ORCA-Fusion BT camera without binning.

**Movie S6.** Live cell imaging movie of three distinct CH12 cells containing a 24xPP7 stem loop array upstream of the endogenous Sα switch region expressing PCP-GFP after 24 hours of stimulation with CIT. A single focal plane was acquired at 6.5 frames per second using a SoRa spinning disc confocal with a 63x objective and a Hamamatsu ORCA-Fusion BT camera without binning.

**Movie S7.** Live cell imaging movie of CH12 cells containing a 24xPP7 stem loop array upstream of the endogenous Sα switch region expressing PCP-GFP after 8 hours of stimulation with CIT. A single focal plane was acquired at 51 frames per second using an Olympus cellTIRF Microscope with a 60x objective and a Hamamatsu ORCA Quest camera at 2x2 binning.

**Movie S8.** Live cell imaging movie of CH12 cells containing a 24xPP7 stem loop array upstream and a 24xMS2 stem loop array downstream of the endogenous Sα switch region expressing PCP-GFP (left panel, green in merge) and MCP-mScarlet (middle panel, magenta in merge) after 8 hours of stimulation with CIT. A Z-stack of the cells was acquired every minute using a SoRa spinning disc confocal with a 63x objective and a Hamamatsu ORCA-Fusion BT camera without binning. The movie is a maximum intensity projection of the Z-stacks.

**Movie S9.** Live cell imaging movie of CH12 cells containing a 24xPP7 stem loop array upstream and a 24xMS2 stem loop array downstream of the endogenous Sα switch region expressing PCP-GFP (green) and and MCP-mScarlet (magenta) after 24 hours of stimulation with CIT. A single focal plane was acquired at 2.5 frames per second alternating 200 ms exposure for the two imaging channels using a SoRa spinning disc confocal with a 63x objective and a Hamamatsu ORCA-Fusion BT camera without binning.

**Movie S10.** Live cell imaging movie of CH12 cells containing a 24xPP7 stem loop array within the first intron of the *AID* locus inserted by RMCE expressing PCP-GFP after 12 hours of stimulation with CIT. A Z-stack of the cells was acquired every minute using a SoRa spinning disc confocal with a 63x objective and a Hamamatsu ORCA-Fusion BT camera without binning. The movie is a maximum intensity projection of the Z-stacks.

**Movie S11.** Live cell imaging movie of CH12 cells containing a 24xPP7 stem loop array upstream of the miniSα switch region inserted by RMCE into the intron of the endogenous *AID* locus expressing PCP-GFP after 8 hours of stimulation with CIT. A Z-stack of the cells was acquired every minute using a SoRa spinning disc confocal with a 63x objective and a Hamamatsu ORCA-Fusion BT camera without binning. The movie is a maximum intensity projection of the Z-stacks.

**Movie S12.** Live cell imaging movie of CH12 cells containing a 24xPP7 stem loop array upstream of the miniSα switch region inserted by RMCE replacing the endogenous Sα region of the *IgH* locus expressing PCP-GFP after 8 hours of stimulation with CIT. A Z-stack of the cells was acquired every minute using a SoRa spinning disc confocal with a 63x objective and a Hamamatsu ORCA-Fusion BT camera without binning. The movie is a maximum intensity projection of the Z-stacks.

**Movie S13.** Live cell imaging movie of CH12 cells containing a 24xPP7 stem loop array inserted by RMCE replacing the endogenous Sα region of the *IgH* locus expressing PCP-GFP after 8 hours of stimulation with CIT. A Z-stack of the cells was acquired every minute using a SoRa spinning disc confocal with a 63x objective and a Hamamatsu ORCA-Fusion BT camera without binning. The movie is a maximum intensity projection of the Z-stacks.

**Movie S14.** Single-molecule imaging movie of CH12 cells expressing AID-HaloTag from the endogenous *AID* locus sparsely labeled with JFX650 HaloTag ligand after 8 hours of stimulation with CIT superimposing single-molecule trajectories generated from single particle tracking analysis of nuclear AID-Halo signals. A single focal plane was acquired at 102 frames per second using an Olympus cellTIRF Microscope with a 60x objective and a Hamamatsu ORCA-Fusion BT camera without binning.

**Movie S15.** Single-molecule imaging movie of CH12 cells transiently expressing the HaloTag protein sparsely labeled with JFX650 HaloTag ligand. A single focal plane was acquired at 102 frames per second using an Olympus cellTIRF Microscope with a 60x objective and a Hamamatsu ORCA-Fusion BT camera without binning.

**Movie S16.** Single-molecule imaging movie of CH12 cells transiently expressing the H2B-HaloTag protein sparsely labeled with JFX650 HaloTag ligand. A single focal plane was acquired was acquired at 102 frames per second using an Olympus cellTIRF Microscope with a 60x objective and a Hamamatsu ORCA-Fusion BT camera without binning.

**Movie S17.** Single-molecule imaging movie of CH12 cells expressing the H56A variant of AID-HaloTag inserted into the endogenous *AID* locus by RMCE sparsely labeled with JFX650 HaloTag ligand after 8 hours of stimulation with CIT. A single focal plane was acquired at 102 frames per second using an Olympus cellTIRF Microscope with a 60x objective and a Hamamatsu ORCA-Fusion BT camera without binning.

**Movie S18.** Single-molecule imaging movie of CH12 cells expressing the E58A variant of AID-HaloTag inserted into the endogenous *AID* locus by RMCE sparsely labeled with JFX650 HaloTag ligand after 8 hours of stimulation with CIT. A single focal plane was acquired at 102 frames per second using an Olympus cellTIRF Microscope with a 60x objective and a Hamamatsu ORCA-Fusion BT camera without binning.

**Movie S19.** Single-molecule imaging movie of CH12 cells expressing the R190X variant of AID-HaloTag inserted into the endogenous *AID* locus by RMCE sparsely labeled with JFX650 HaloTag ligand after 8 hours of stimulation with CIT. A single focal plane was acquired at 102 frames per second using an Olympus cellTIRF Microscope with a 60x objective and a Hamamatsu ORCA-Fusion BT camera without binning.

**Movie S20.** Single-molecule imaging movie of CH12 cells in which all switch regions were deleted expressing AID-HaloTag from the endogenous *AID* locus sparsely labeled with JFX650 HaloTag ligand after 8 hours of stimulation with CIT. A single focal plane was acquired at 102 frames per second using an Olympus cellTIRF Microscope with a 60x objective and a Hamamatsu ORCA-Fusion BT camera without binning.

**Movie S21.** Dual color imaging of CH12 cells containing a 24xPP7 stem loop array upstream of the endogenous Sα switch region expressing AID-HaloTag from the endogenous *AID* locus sparsely labeled with JFX650 HaloTag ligand (middle panel, magenta in merge) and PCP-GFP (left panel, green in merge) after 24 hours of stimulation with CIT. Both channels were acquired simultaneously at 1 frame per second using two Hamamatsu ORCA-Fusion BT cameras on a Olympus cellTIRF Microscope with a 60x objective.

---

**Supplemental Figure 1.**
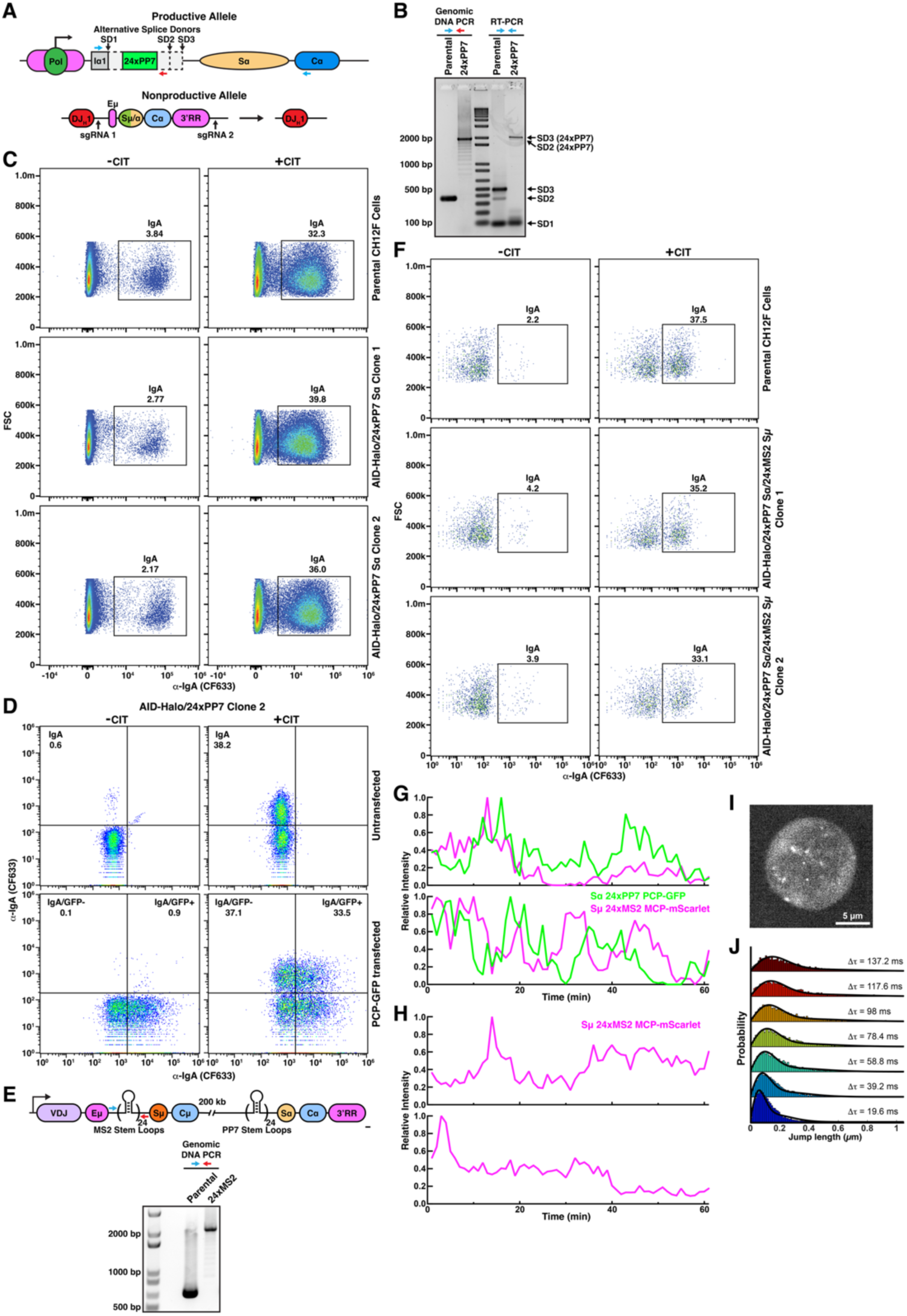
**(A)** Model of the two *IgH* alleles in the engineered CH12 cell line containing a 24xPP7 stem loop array in the productive *IgH* locus allele. **(B)** Genomic DNA PCR (left) and RT-PCR (right) using the primer locations indicated in (A). **(C)** CSR assays of parental CH12 cells and CH12 cell lines containing the 24xPP7 stem loop array upstream of the Sα switch region and a HaloTag at the C-terminus of the endogenous AID coding region. **(D)** CSR assays of CH12 cells containing a 24xPP7 stem loop array upstream of the Sα switch region and a HaloTag at the C-terminus of the endogenous AID coding region transfected with a plasmid encoding PCP-GFP. **(E)** Genomic DNA PCR using the indicated primers to confirm 24xMS2 stem loop array insertion upstream of the Sµ switch region. **(F)** CSR assays of parental CH12 cells and CH12 cell lines containing a 24xPP7 stem loop array upstream of the Sα switch region, a 24xMS2 stem loop array upstream of the Sµ switch region, and a HaloTag at the C-terminus of the endogenous AID coding region. **(G-H)** Quantification of **(G)** co-localized static PCP-GFP and MCP-mScarlet signals, or **(H)** MCP-mScarlet signal without the presence of PCP-GFP signal in CH12 cells containing a 24xMS2 stem loop array upstream of the Sµ switch region and a 24xPP7 stem loop array upstream of the Sα switch region. **(I)** Image of a CH12 cell containing a 24xPP7 stem loop array upstream of the Sα switch region expressing PCP-GFP without an NLS 8 hours after stimulation with CIT. **(J)** Step size distributions and two state fits of single-particle trajectories generated from single-molecule imaging of the PCP-GFP labeled Sα germline transcript.

**Supplemental Figure 2.**
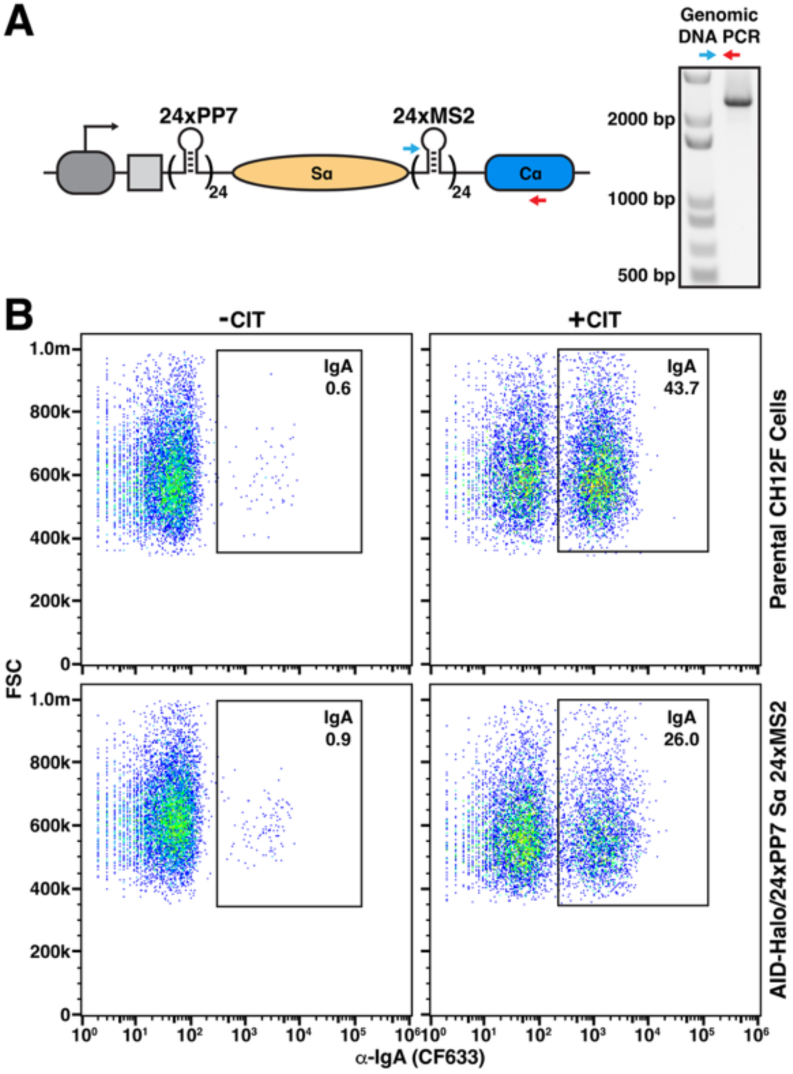
**(A)** PCR of genomic DNA using the indicated primer pair confirming the insertion of the 24xMS2 stem loop array downstream of the Sα switch region. **(B)** CSR assays of parental CH12 cells and CH12 cell lines containing a 24xPP7 stem loop array upstream of the Sα switch region, a 24xMS2 stem loop array downstream of the Sα switch region, and a HaloTag at the C-terminus of the endogenous AID coding region.

**Supplemental Figure 3.**
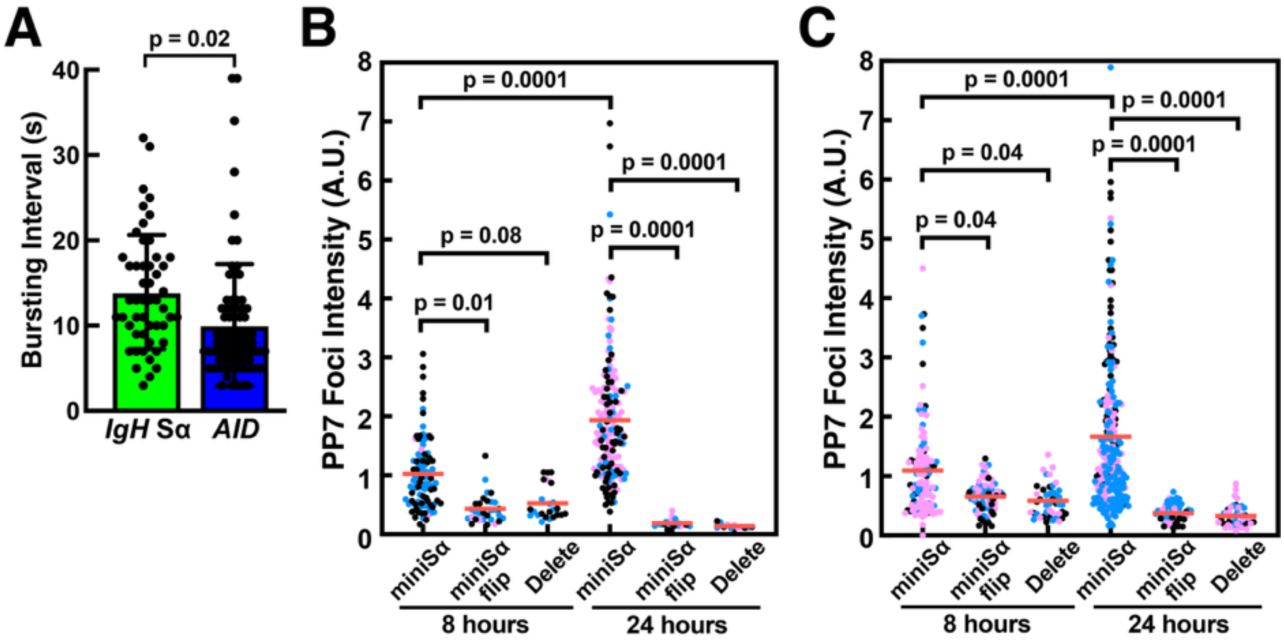
**(A)** Quantification of the transcriptional burst interval of CH12 cells containing a 24xPP7 stem loop array in the *IgH* locus Sα switch region or intron 1 of the *AID* locus (N = 22 and 25 cells, Mean±SD, T-Test). **(B-C)** Quantification of the intensity of nascent RNA signals marked by PCP-GFP in CH12 cells with **(B)** an engineered Sα switch region in the *IgH* locus or **(C)** an engineered *AID* locus containing only a 24xPP7 stem loop array (delete), or a 24xPP7 stem loop array with either the miniSα sequence or the reverse complement of the miniSα sequence (flip) (N = 3 biological replicates, n > 10 cells per replicate, two-tailed T-Test).

**Supplemental Figure 4.**
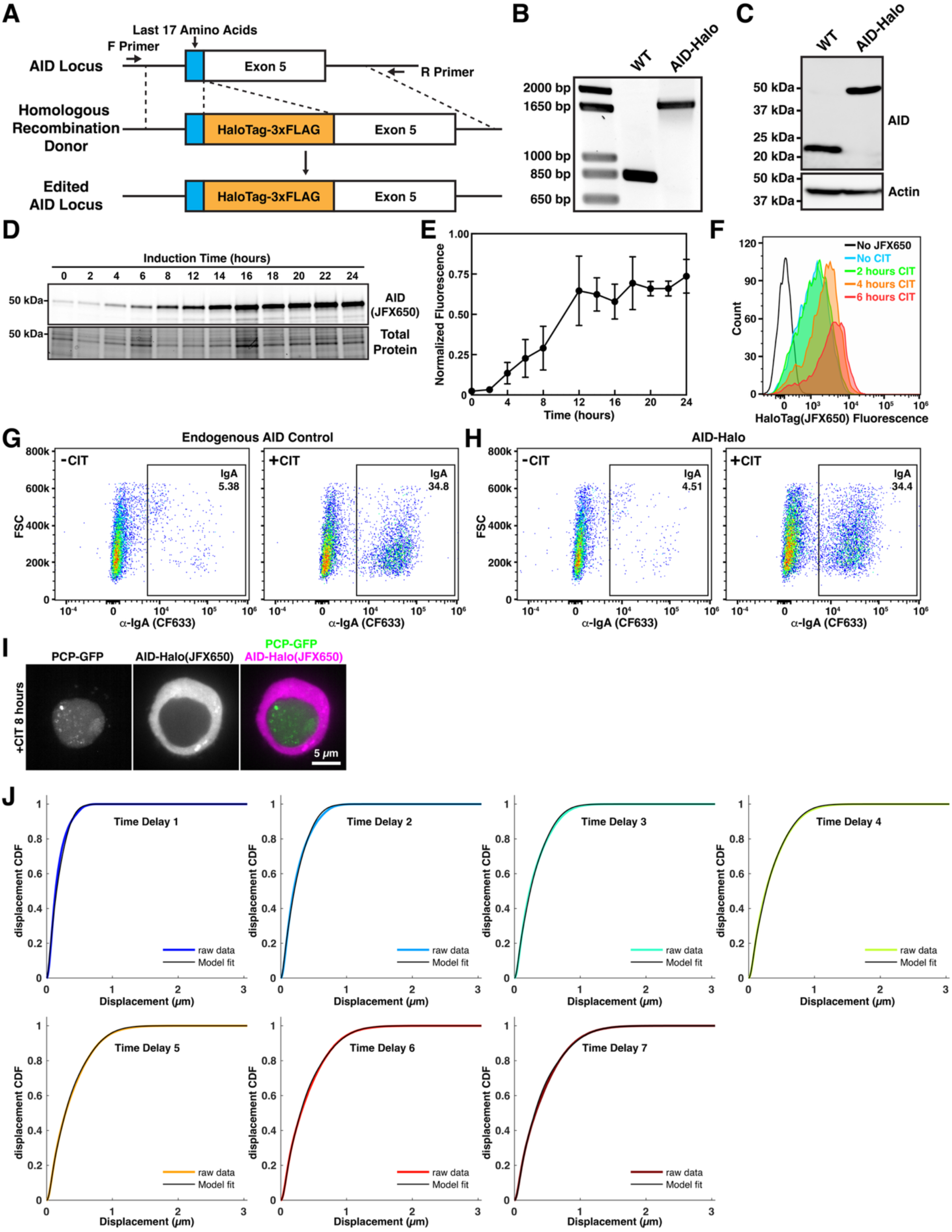
Functional characterization of HaloTagged AID expressed from the endogenous *AID* locus. **(A)** Genome editing approach to insert the Halo-3xFLAG tag at the C-terminus of the AID coding sequence of the endogenous *AID* locus. **(B)** Agarose gel of PCR products amplified from the endogenous *AID* locus from control cells and cells containing the coding sequence for the Halo-3xFLAG tag. **(C)** Western blot of cell lysates from control cells and CH12 cells expressing AID-Halo from the endogenous *AID* locus probed with antibodies targeting AID and actin. **(D)** Fluorescence imaging (top) and total protein staining (bottom) of SDS-PAGE gel loaded with cell lysates generated from cells stained with the fluorescent JFX650 HaloTag-ligand after induction of CSR using CIT for the indicated time. **(E)** Quantification of AID expression level over time using the fluorescence intensity of AID-Halo stained with JFX650 HaloTag-ligand (N = 3 biological replicates, Mean±SD). **(F)** Flow cytometry analysis of the expression of AID-Halo stained with JFX650 HaloTag-ligand over time. **(G-H)** Analysis of CSR efficiency using flow cytometry with an antibody targeting cell surface localized IgA antibodies in **(G)** control and **(H)** CH12 cell expressing AID-Halo from the endogenous *AID* locus. **(I)** Image of a living CH12 cell containing a 24xPP7 stem loop array upstream of the Sα switch region expressing PCP-GFP, and AID-Halo from the endogenous *AID* locus quantitatively stained with JFX650 HaloTag-ligand. **(J)** Spot-On fitting using a 3-State model of cumulative density function of Halo-AID displacements generated from single-particle tracking of nuclear AID trajectories 8 hours after CIT induction.

**Supplemental Figure 5.**
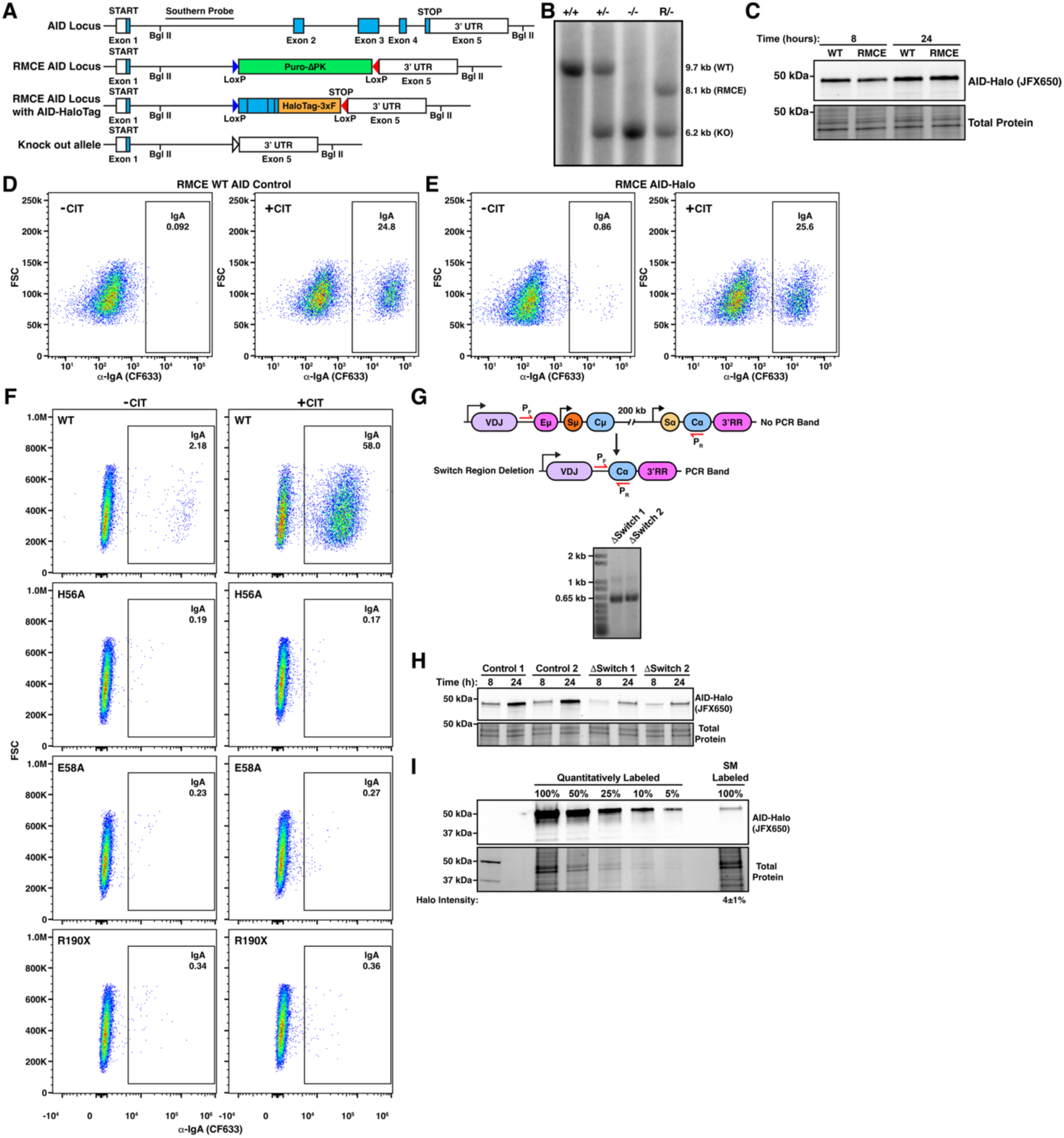
**(A)** Model of the endogenous *AID* locus and different variants of the endogenous AID locus containing an RMCE cassette. **(B)** Southern blot using the probe location indicated in (A) of CH12 cells engineered to contain an RMCE cassette in one *AID* allele and a deletion in the second *AID* allele. **(C)** Fluorescence gel of cell lysates labeled with JFX650 HaloTag-ligand generated from CH12 cells expressing AID-Halo by direct insertion into the *AID* locus (WT) or using RMCE to insert the HaloTag into the *AID* locus. **(D-E)** CSR assays of **(D)** CH12 cells expressing AID-Halo by direct insertion into the *AID* locus (WT) or **(E)** using RMCE to insert the HaloTag into the *AID* locus. **(F)** CSR assays of CH12 cells expressing different variants of AID-Halo inserted into the *AID* locus using RMCE. **(G)** Experimental design and PCR analysis of deletion of all switch regions from the IgH locus in CH12 cells expressing AID-Halo from the endogenous *AID* locus. **(H)** Fluorescent gel (top) and total protein stain (bottom) of SDS-PAGE gel loaded with cell lysates generated from control CH12 cells and CH12 cells lacking the switch region sequences expressing AID-Halo stained with JFX650 HaloTag-ligand. **(I)** Fluorescent gel of cell lysates from CH12 cells expressing AID-Halo from its endogenous locus. The HaloTag was either quantitatively labeled (100 nM JFX650 HaloTag-ligand for 30 minutes) or labeled for single-molecule imaging (0.6 nM JFX650 HaloTag-ligand for 30 seconds). Quantification (below) from 3 biological replicates (Mean±SD).

